# Chain of Structures: Connecting Reasoning and Generation for Protein Design

**DOI:** 10.64898/2026.10.09.758133

**Authors:** Qingran Cai, Mingyang Xu, Jiaxian Yan, Zhicai Wang, Yufeng Liu, Qizhi Pei, Zaixi Zhang

## Abstract

Scientific design requires reasoning in scientific spaces that represent the objects being designed. Building on Scinetics’ *Chain of Science Tokens*, we present *chain-of-structures* (COS) as a concrete protein-design case study that connects reasoning and generation through explicit structural states. In this technical report, we apply COS to enzyme design, where catalytic function requires precise local geometry within a compatible global fold, making it a demanding test of reasoning from chemical requirements to protein structure. Starting from a reaction specification and target length, COS couples a language reasoning expert with an all-atom generation expert across **four rounds: catalytic core, active site, global backbone, and complete sequence and structure**. Each reasoning state directly conditions generation, and the resulting structure becomes context for the next decision. Discrete residue constraints and continuous latents connect the two experts, enabling structural supervision of the reasoning interface. On the evaluated benchmark, COS leads the compared baselines on five of six metrics, achieving 57.0% EC3 agreement and a TM-score of 0.729.

## 1 From Science Tokens to Protein Design

### Reasoning through scientific states

Scinetics’ multimodal AI4S program aims to connect scientific understanding, reasoning, and generation within a common modeling framework. *Chain of Science Tokens* expresses this goal as a sequence of typed scientific states: language specifies a goal, scientific representations express intermediate decisions, and generation produces a candidate that can be inspected and refined. In this report, science tokens encompass residue identities and structured molecular representations, including continuous latents. *chain-of-structures* provides a concrete case study of this principle in protein design. The implementation reported here is an exploratory prototype, not the final model of Scinetics’ multimodal AI4S program. A universal discrete vocabulary across scientific modalities remains a broader research direction.

### Why enzymes?

An enzyme must organize a chemical reaction. Catalytic side chains, substrate orientation, hydrogen bonding, electrostatics, and the surrounding scaffold jointly shape its activity. A plausible global fold alone is insufficient: small local changes can disrupt the arrangement required for catalysis. Enzymes therefore expose whether a design process can connect chemical intent to local geometry and then to a complete protein [1, 2, 3].

### Connecting reasoning and generation

Protein generators, inverse-folding models, and multimodal protein models supply complementary capabilities [4, 5, 6, 7]; GENzyme constructs reaction-conditioned pockets before full enzymes [8]. Proteo-R1 couples a molecular understanding expert to a diffusion generator through residue-level commitments [9]. ProteinReasoner uses evolutionary profiles as intermediate reasoning representations linking protein structure and sequence [10]. SciReasoner makes structural tokens explicit evidence for reasoning across proteins, molecules, and crystals [11]. Building on these directions, *chain-of-structures* repeatedly connects chemical reasoning to structural generation through four rounds. Each generated structure informs the next decision, while a continuous interface carries structural training signals back to the reasoner.

## 2 A Four-Round Chain of Structures

The chain follows a chemical-to-structural hierarchy. Each round pairs a reasoning record with the part of the design it commits. **The mechanistic premise carries no protein coordinates and is not a round of its own; it opens Round 1. Rounds 1–4 then progressively specify the molecule**, so four scientific rounds are four structure-bearing updates.

A design record is written one round at a time and is the only thing that passes between the two experts. Round *k* produces *r*_*k*_ = (prose, *π*_*k*_), where the prose states why this part of the molecule must look the way it does and *π*_*k*_ = { (*p, a*_*p*_) } is a set of (canvas position, residue) pairs: *p* ∈ {1, …, *L*} on a canvas of the requested length *L*, and *a*_*p*_ one of the twenty residues. Positions are absolute rather than relative because the generator is conditioned position-wise; a pair is therefore directly executable, and the prose beside it is what makes the commitment auditable. The record is append-only: Round *k* may add pairs but never edits *π*_1_, …, *π*_*k−*1_, so the chain is a monotone refinement rather than a search that backtracks. Section 6 returns to what backtracking would require.

### 2.1 From reaction constraints to a complete protein

Figure 1 develops one illustrative FAD synthetase design: the reaction constrains the relative orientation of the reactive phosphates, the catalytic core then constrains substrate recognition, the site constrains the backbone, and the backbone constrains sequence completion. Each round explains why its commitment follows from the preceding one. The specification here is an explicit reaction; a substrate alone would only induce a hypothesis about function, since pinning a unique transformation generally needs a reaction or a product. The record is both an interpretable object and a conditioning interface, and its chemical hypotheses still require evaluation.

**Figure 1.**
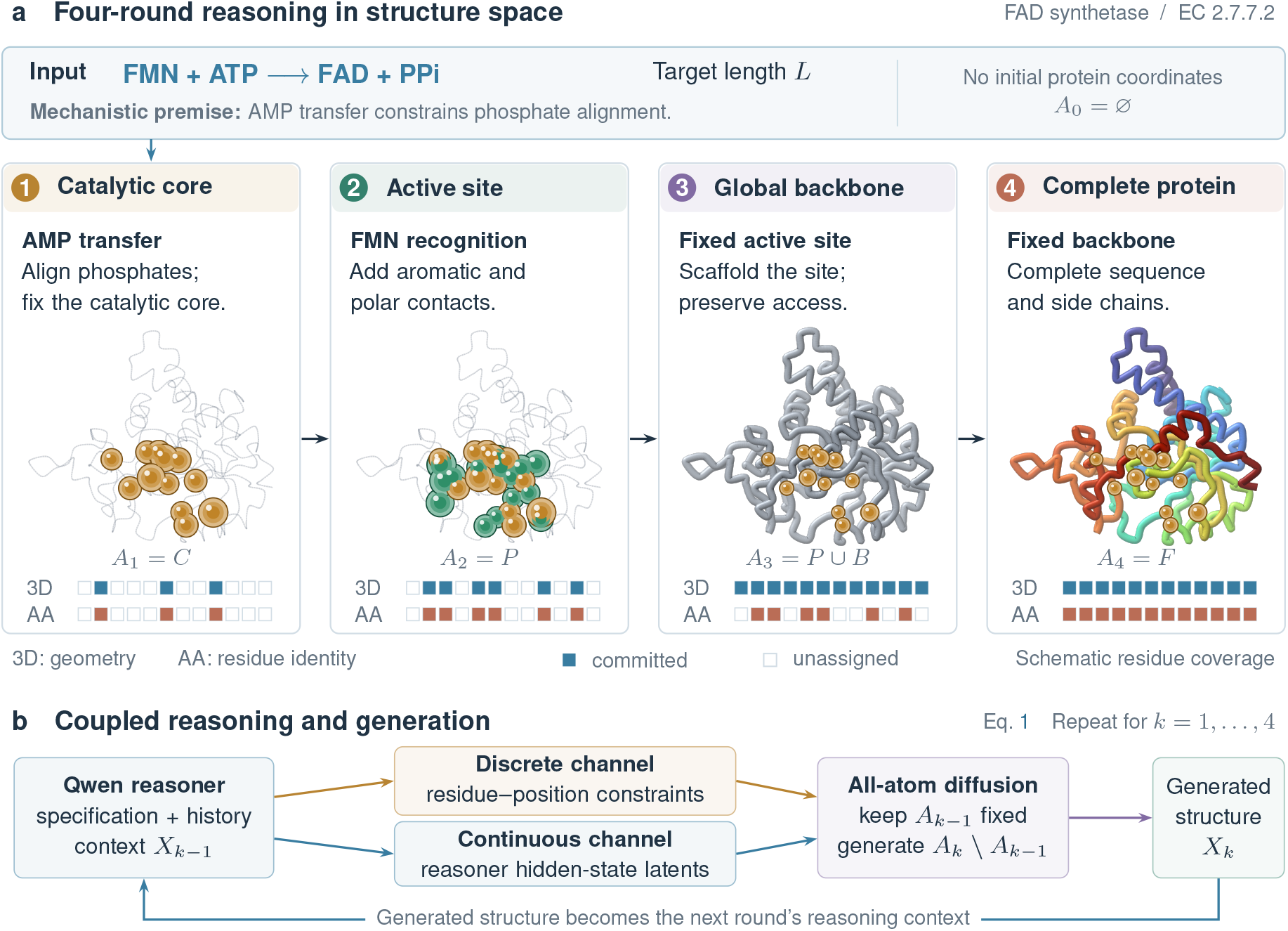
*chain-of-structures*: a four-round FAD synthetase design example. Panel a expands the reaction FMN + ATP→FAD + PP_*i*_ into the mechanistic constraints Round 1 opens from, then the four committed rounds: catalytic-core geometry, substrate recognition, a supporting backbone, and sequence completion. The worked design example follows the chemistry and flavin-recognition logic of FMN adenylyltransferase [12]. The 3D and AA strips show committed geometry and amino-acid information. Panel b connects the reasoner to the diffusion head through residue identities and continuous latents, then returns the generated structure as context. Structural illustrations are retained from the earlier manuscript.

### 2.2 Four-round inference

Inference consumes a reaction and a target length and runs four rounds. At round *k* the reasoner reads the specification *u*, the records *r*_*<k*_ it has already written, a shortlist *S*_*k*_ of candidate positions, and the structure *X*_*k−*1_ the generator produced in the previous round; it emits *r*_*k*_. The generator is then re-conditioned on *π*_*≤k*_, asked to commit *A*_*k*_ while holding *A*_*k−*1_ fixed, and returns *X*_*k*_, which becomes the next round’s structural context. Round 1 is the one round with no structural context: it argues from the mechanistic premise and *S*_1_ alone. Each structural update is a full 200-step sampling run, not a partial refinement of the previous sample, so a round’s commitments are realized rather than nudged. LigandMPNN redesign is an explicitly separate sequence-evaluation branch applied after structure generation and is never part of the chain.

#### The shortlist

*S*_*k*_ turns “name positions on a blank canvas” into “choose among candidates”, which is the difference between an unconstrained and a well-posed decision. It holds 100 rows. A row names a canvas position and carries five fields for it: the residue that the aligned training enzyme places there, how far that alignment anchor is, which tenth of the chain the position falls in, the residue most often catalytic in that tenth for this reaction class, and the position’s overall prior rank. Rows are ordered by position, not by score, so the reasoner cannot read the ranking off the row order. *S*_*k*_ is built by lookup and contains nothing measured on the target protein: positions are ranked by a class-conditioned residue-position prior and by alignment to the catalytic site of the closest-length training enzyme of the same class. The target’s own reaction is held out of that library, so a near-duplicate cannot be copied; without that exclusion the shortlist nearly contains the answer, which is why we report the lookup-only ranking as a baseline rather than folding it into the method. A round supervises the positions of its state that *S*_*k*_ contains and records that coverage: a position outside *S*_*k*_ is one the reasoner could not have chosen at inference either, so counting it as an error would measure the shortlist rather than the reasoner.

## 3 Learning and Executing the Chain

### 3.1 Two experts and a dual-channel interface

The current implementation uses Qwen as the reasoning expert and an RFdiffusion3-based all-atom *diffusion head* for generation [13, 5]. Following the reasoning-to-generation interface of Proteo-R1 [9], the reasoner returns, for every position it commits in round *k*, a discrete residue identity 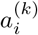 and a continuous latent 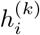 taken from the reasoner’s hidden state for that commitment. Both enter the generator where its own conditioning already enters, so no new pathway into the denoiser is introduced:

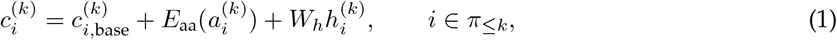

where 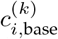 is the generator’s own per-residue conditioning for position *i* (the representation it would use with no record at all, carrying the ligand context and whatever structure is already pinned), *E*_aa_ is the generator’s existing residue-type embedding, and *W*_*h*_ is a learned linear map from the language model’s hidden width into the generator’s conditioning width. Positions outside *π*_*≤k*_ keep 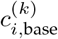 unchanged, so a round writes only where it has something to say and the rest of the canvas is left to the generator’s prior.

Two properties of Equation 1 matter. First, the two terms carry different information and are supervised differently: the residue identity is a discrete decision that passes no gradient, while 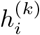 is continuous and does, so structural error reaches the reasoner only through the latent. The identity term is what makes a commitment legible; the latent term is what makes it trainable. Second, *W*_*h*_ is zero-initialized, so at the start of joint training the interface is exactly the identity on *c*_*i*,base_ and the generator behaves as the pretrained model it is. The reasoner has to earn its influence rather than be granted it, which keeps an untrained interface from destroying a capable generator early in training. End-to-end here means training through this continuous interface; it does not mean differentiating through the discrete choice, nor through the full inference-time sampling trajectory.

### 3.2 A shared four-round schedule

Let *C* be catalytic-core atoms, *P* ⊇ *C* the active-site atoms, *B* full-chain backbone atoms, and *F* ⊇ *P* ∪ *B* the complete protein atom set. Ligand atoms remain conditioning context. Let *A*_*k*_ denote protein atoms committed *after* Round *k* (*A*_0_ is the empty protein state):

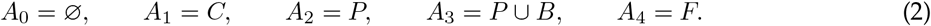

These sets are nested by construction, which is what lets one generator serve all four rounds: round *k* is the same conditional sampling problem with a larger given set and a larger target. At update *k* the generator is given the atoms in *A*_*k−*1_, taken from the structure it produced in round *k* − 1 rather than from the reference, and is asked for *A*_*k*_\*A*_*k−*1_; atoms outside *A*_*k*_ remain unconstrained. Supervision is applied on *A*_*k*_ only. Keeping the given set and the supervised set disjoint is what prevents a round’s target from being described as something the generator was already handed, which would make the loss measure copying rather than design. Because the given set is a *generated* structure at inference and a generated structure during Phase III as well, the train-time and test-time conditioning distributions match; conditioning on reference atoms would make every round easier than it is at deployment.

### 3.3 Training: alignment, reasoning, and joint design

The training objective pairs the four reasoning records with the four structural losses:

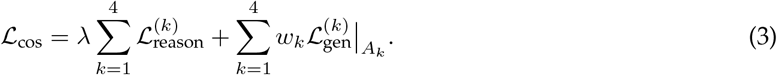

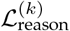is the token-level cross-entropy of the record *r*_*k*_ under the reasoner, conditioned on the specification, the earlier records and the generated context of round *k*. The prose and the committed pairs are one sequence, so the same loss trains the explanation and the decision. The pair tokens are a small fraction of that sequence, so they are up-weighted; without that the objective is dominated by prose it already predicts well. 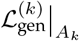 is the generator’s denoising loss at noise levels sampled as in pretraining, restricted to the atoms of *A*_*k*_, with the atoms of *A*_*k−*1_ given. The per-round weights *w*_*k*_ and the balance *λ* are the two knobs that decide whether the reasoner is trained to be articulate or to be executable; we set them so that no single round dominates the structural term, since the rounds differ by more than an order of magnitude in how many atoms they supervise. Starting from pretrained experts, we complete post-training in **one week on eight NVIDIA H100 GPUs**. The three phases build complementary capabilities: reading molecular inputs, reasoning about structural constraints, and converting those decisions into generated structures. Each one trains a different, disjoint set of parameters while the rest stay frozen. They are distinct from the four design rounds executed for each enzyme: the phases are stages of training, the rounds are stages of a single design. Staging the *parameters* rather than the objective is deliberate; a schedule that instead switched losses on and off was what degraded the reasoner’s ability to read its own inputs in an earlier iteration.

#### Phase I: multimodal alignment

We train the input projector while keeping Qwen and the diffusion expert frozen. Ligand-atom schema and caption supervision align molecular features with the language model’s input space. This phase establishes a readable molecular interface while preserving the pretrained experts, so later reasoning can use molecular context through a calibrated representation.

#### Phase II: structure-grounded reasoning

With the structural encoder fixed, we train Qwen adapters and auxiliary heads on per-residue structural labels and four-round reasoning records. Secondary structure, solvent accessibility, ligand contacts, distances, and pocket membership teach the model to read local and global geometry; expert-guided traces teach it to turn these observations into ordered design decisions. Replay helps retain earlier capabilities. This phase prepares the reasoner to specify meaningful constraints before joint design.

#### Phase III: joint reasoning-guided generation

We optimize LoRA adapters in both experts and the connecting interface using the reasoning and structural losses in Equation 3. Structural supervision is denoising at sampled noise levels, with core, site, backbone, and full-protein targets introduced progressively. Gradients through the continuous channel teach the reasoner to produce conditioning that the diffusion head can realize. This final phase connects the scientific quality of a decision to its geometric consequences.

## 4 Four-Round Data and Supervision

The four-round schema organizes each record as a specification and four aligned rounds:

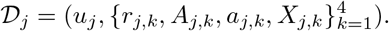

Here *r*_*j,k*_ is the reasoning target, *A*_*j,k*_ the committed atom set, *a*_*j,k*_ the available residue identities, and *X*_*j,k*_ its geometry. The schema aligns each reasoning round with its structural commitment and makes the final sequence decision explicit.

Figure 2 shows the construction end to end; the subsections below give the definitions and the measurements behind each step.

**Figure 2.**
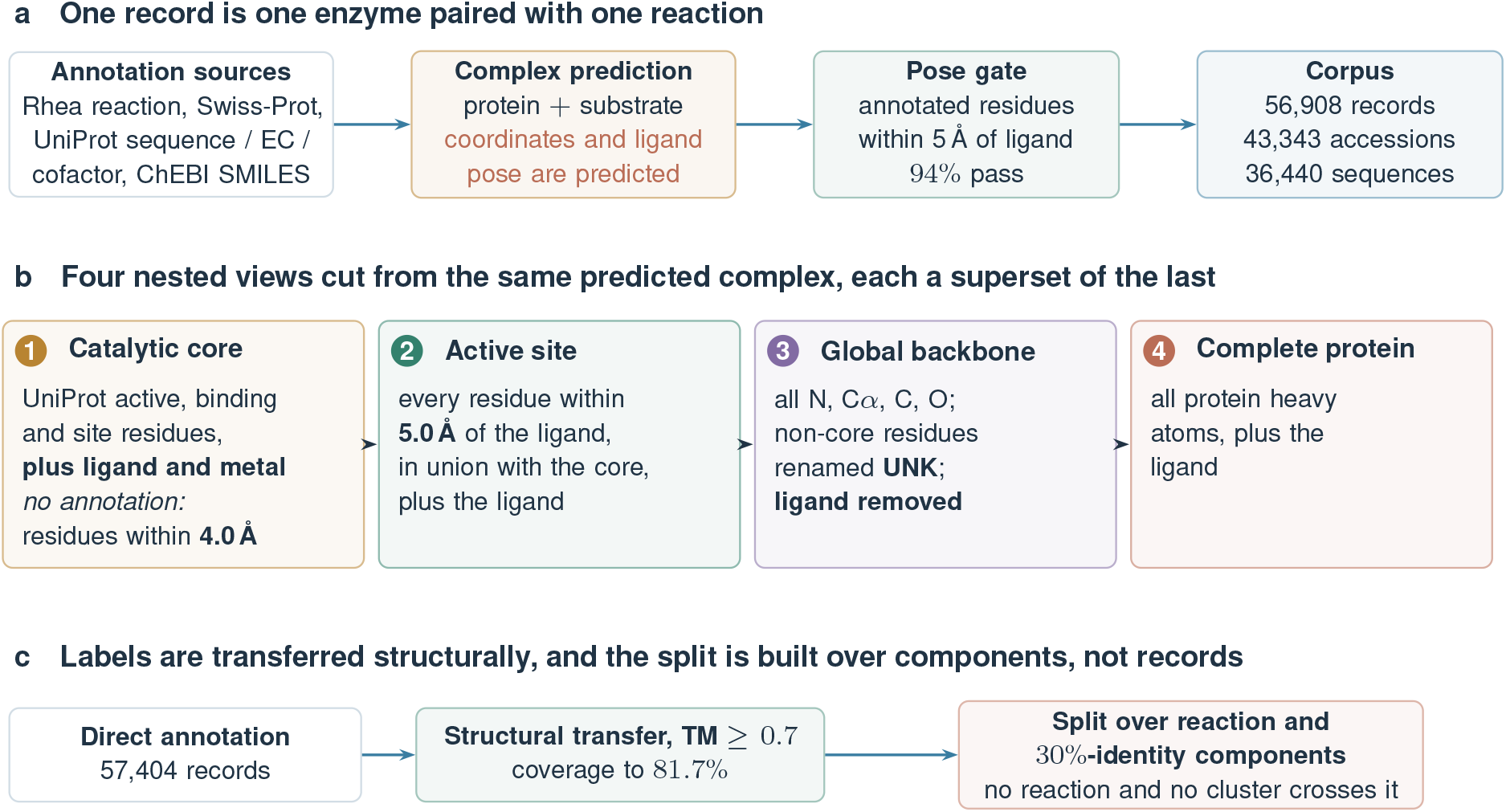
How a four-round record is built. Panel a: records are assembled from annotation databases and a predicted protein-ligand complex, and a pose gate removes predictions that place the ligand outside the annotated site. Panel b: the four structural targets, with the cutoffs exactly as computed. Panel c: catalytic-site labels are extended by structural transfer above a calibrated similarity threshold, and the split is assigned over connected components so that neither a reaction nor a sequence cluster can straddle it.

### 4.1 Where the records come from

Every record is one enzyme paired with one reaction. The corpus is assembled rather than downloaded. Rhea reaction entries are joined to Swiss-Prot through the Rhea-to-UniProt mapping, UniProt supplies the sequence, the four-level EC number and the cofactor annotation, and Rhea and ChEBI supply the substrate as SMILES. Each protein and its substrate are written into a complex-prediction job, and the predicted complex is kept as the record’s structure. The upstream filter removes entries without an EC number, entries whose substrate equals their product, EC class 7, electron-transport entries, and chains longer than 1,000 residues. The resulting corpus holds **56**,**908 records** over **43**,**343 UniProt accessions** and **36**,**440 distinct sequences**.

#### The structures are predictions, and we say so

The coordinates and the ligand pose are both predicted, not experimental. This is a property of the domain rather than a shortcut: of 600 sampled training accessions, only 5% have any experimental structure in the PDB, and holo structures with the substrate bound are rarer still. For 95% of these enzymes an experimental structure does not exist. Catalytic-residue labels are likewise mostly inferred rather than observed. Across UniProt, roughly 107,646 proteins carry an activesite annotation, but only about 1,500 to 2,400 carry one with direct experimental evidence; the remainder come from rule-based inference or homology transfer. We therefore treat both the geometry and the site labels as predicted-tier supervision throughout, and we do not describe any structural agreement in this paper as agreement with experiment.

#### Pose quality is checked before a record is used

A predicted complex is only useful here if the ligand sits in the annotated site, because every structural target below is defined relative to that ligand. We measure, per record, the fraction of annotated active-site residues lying within 5 Å of the predicted ligand. Over a 120-record sample the mean is 0.97 and the median is 1.00: 94% of records place at least four fifths of the annotated residues in contact, 3% are partial, and 3% misplace the ligand. Records in the last group are dropped, since a misplaced ligand corrupts the core and the pocket together. Combined with the 79% of records that carry a catalytic annotation at all, this leaves rather more than half the corpus usable for four-round supervision.

### 4.2 Four nested structural targets

The four rounds are supervised by four nested views of the same predicted complex, each a strict superset of the previous one. The definitions are mechanical and are given here exactly as they are computed, because the choice of cutoff decides what each round is actually asked to produce.

1. **Catalytic core**. The UniProt active-site, binding-site and other-site residues that map into the structure, together with the ligand and any metal. When a record carries no such annotation, the core falls back to the residues whose heavy atoms lie within 4.0 Å of the ligand.
2. **Active site**. Every protein residue with a heavy atom within 5.0 Å of the ligand, in union with the core, again together with the ligand. The 5 Å shell is what makes this round a recognition problem rather than a repetition of the core.
3. **Global backbone**. All protein residues reduced to their N, C*α*, C and O atoms, with every residue outside the core renamed to UNK and the ligand removed. The round therefore receives a scaffold that still pins the catalytic identities but discloses no other side chain and no substrate.
4. **Complete protein**. All protein heavy atoms together with the ligand.

Because the four views are cut from one predicted complex, they share a coordinate frame exactly. Backbone and full-complex C*α* positions agree to within 0.0005 Å with no alignment step, and one residue and atom mapping connects all four views, including active-site residues that are far apart in sequence.

### 4.3 Site labels and their coverage

A four-round record needs to know which residues are catalytic, and the annotation does not cover the corpus. Direct UniProt active-site annotations reach 57,404 of the enzyme records. Transferring labels between structurally similar enzymes raises coverage to **81.7%**, and the transfer has to be structural rather than sequence-based: within a third-level EC class, sequence alignment agrees with the gold label only 41.1% of the time, while structural alignment agrees 82.7% of the time, because enzymes that share a reaction often share a fold without sharing a sequence.

Transfer accuracy depends sharply on structural similarity, so we calibrated it before using it. Below a TM-score of 0.7 the transferred label is right between 2% and 12% of the time and is worthless; between 0.7 and 0.8 it is right 68.6% of the time; above 0.9 it is right every time in our sample. We transfer only above TM 0.7. The remaining 26,917 records have no labelled donor at any threshold: searching the labelled set for each of them returns a best TM-score with median 0.219, and only 0.2% reach 0.7. Their folds are absent from the labelled set, so more searching would not help, and they are excluded rather than labelled by a weaker rule.

### 4.4 Deduplication and the split

Records are not proteins. Of the 56,908 records, 6,911 accessions appear more than once, which accounts for 18,762 records, a third of the corpus, and one accession appears 42 times. Within such a group the sequence, EC list and length are identical and only the coordinates differ, with a median pairwise TM-score of 0.9952 against 0.3249 for random pairs. These are repeated prediction runs of one protein, not distinct structures, and counting them as independent records inflates any sample size computed from them.

The split must therefore be built over proteins and reactions rather than over records. A naive random split is not an honest test here, because only 1,765 distinct Rhea reactions underlie the corpus, so roughly thirteen proteins share each reaction and a randomly chosen test reaction has almost certainly been seen in training. We instead form a graph whose edges join records that share a reaction or fall in the same MMseqs2 cluster at 30% identity, which yields 1,221 clusters and 642 connected components, and assign whole components to one side of the split. The result is verified on the written files: no reaction and no cluster crosses a split boundary.

One property we do not claim is a temporal split. These entries are mostly from around 2008, with a training median of 2008-02-26 against a test median of 2008-05-20, so a modern cutoff date would leave almost nothing. We report novelty with respect to reaction and sequence instead, which is the axis that matters for this task.

The construction contract distinguishes *plans*, based on the specification and previous state, from *observations* of the state just built. For example, a Round 1 plan cannot rely on the global fold introduced at Round 3. Ground-truth structures supply training targets; at inference, the reasoner instead receives the generated structural context from preceding rounds.

#### Four-round supervision example: a serine-hydrolase design

1. **Catalytic core:** hydrolysis motivates a nucleophile, a proton-transfer partner, and stabilization of the tetrahedral intermediate; propose a Ser–His–Asp arrangement and generate its geometry.
2. **Active site:** place supporting groups around the core, including potential oxyanion stabilization and substrate contacts.
3. **Global backbone:** scaffold the committed site with a compatible backbone and access to the substrate-binding region.
4. **Sequence and structure:** complete the surrounding residues and side chains, retain the catalytic commitments, and assess folding compatibility.

### 4.5 Corpus and evaluation scope

The supplied split summary accounts for **22**,**201 records**: **19**,**927 training, 1**,**125 validation**, and **1**,**149 test**; the test pool spans 98 EC classes. These are record-level partitions. The source also lists 22,948 corpus records, leaving 747 outside the provided split accounting; those records are not added to any split or evaluation denominator here. Sequence-cluster separation and the precise length-filter manifest remain necessary for stronger generalization claims.

## 5 Results and Baseline Comparisons

Table 3 presents the **main results**, comparing *chain-of-structures* with four baseline generators in functional matching, structural quality, and sequence-distribution fidelity. Every row is text-to-protein, taking a functional specification as input and producing a sequence and a structure, so the column values are comparable even though the methods differ in what they condition on and how.

**Table 1.** One four-round definition for reasoning, structural supervision, and inference.

| Round | Scientific state | Coordinates committed | Residue identities committed |
| --- | --- | --- | --- |
| n/a | Mechanistic premise (opens Round 1) | None | None |
| 1 | Catalytic core | Catalytic atoms $C$ | Catalytic residues |
| 2 | Active site | Core and support site $P$ | Active-site residues |
| 3 | Global backbone | $P \cup B$ | Active-site residues retained |
| 4 | Sequence + structure | Complete protein $F$ | Full sequence |

**Table 2.** Four-round record construction. Structural size summaries are retained from the current report’s 300-record sample; they are not new full-corpus estimates.

| Round | Reasoning target | Structural / sequence target | Reported size |
| --- | --- | --- | --- |
| n/a | Transformation and required chemical roles | Specification; empty protein state | No protein atoms |
| 1 | Catalytic residue choice and arrangement | Annotated catalytic motif | Median 4 residues |
| 2 | Substrate contacts and local support | Core plus active-site shell | Median 13 residues; 8 sequence islands |
| 3 | Scaffold around the committed site | Full N/Cα/C/O backbone plus retained site | Median 349 residues |
| 4 | Sequence compatibility and side-chain completion | Full sequence and all-atom complex | Complete record |

**Table 3.**
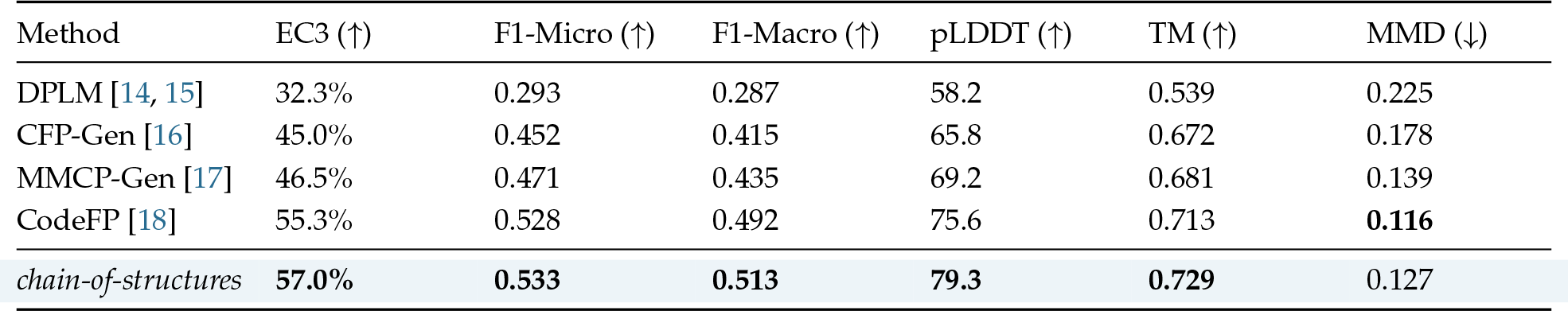
Main results: overall functional matching and structural quality. Comparison of *chain-of-structures* with four baseline generators on the same evaluation records using identical scorers. EC3 (third-level EC agreement under CLEAN) and micro-/macro-F1 assess predicted functional matching; ESMFold pLDDT and TM summarize structural confidence and self-consistency; MMD measures deviation from natural *k*-mer spectra. Baselines are retrained on the same training split, each row reporting the longest run we trained. We do not claim any row is at its ceiling: baseline EC3 keeps moving with training budget in our own measurements, so these are budget-matched points rather than converged optima. Arrows indicate the preferred direction; **bold** marks the best value in each metric column.

| Method | EC3 (↑) | F1-Micro (↑) | F1-Macro (↑) | pLDDT (↑) | TM (↑) | MMD (↓) |
| --- | --- | --- | --- | --- | --- | --- |
| DPLM [14, 15] | 32.3% | 0.293 | 0.287 | 58.2 | 0.539 | 0.225 |
| CFP-Gen [16] | 45.0% | 0.452 | 0.415 | 65.8 | 0.672 | 0.178 |
| MMCP-Gen [17] | 46.5% | 0.471 | 0.435 | 69.2 | 0.681 | 0.139 |
| CodeFP [18] | 55.3% | 0.528 | 0.492 | 75.6 | 0.713 | **0.116** |
| *chain-of-structures* | **57.0%** | **0.533** | **0.513** | **79.3** | **0.729** | 0.127 |

**Table 4.** Ablations: what each half of the interface is worth. Same records, same scorers, and same sampling budget as Table 3. *No chain of reasoning* removes the written rounds and conditions the generator on the specification alone, leaving the continuous channel in place. *No continuous channel* keeps the written rounds but injects only the discrete residue– position pairs, so structural supervision can no longer reach the reasoner. Arrows give the direction of improvement; ∆ is EC3 relative to the full system.

| Variant | EC3 (↑) | F1-Micro (↑) | F1-Macro (↑) | pLDDT (↑) | TM (↑) | MMD (↓) | ΔEC3 |
| --- | --- | --- | --- | --- | --- | --- | --- |
| *chain-of-structures* (full) | **57.0%** | **0.533** | **0.513** | **79.3** | **0.729** | **0.127** | n/a |
| no chain of reasoning | 53.4% | 0.491 | 0.481 | 72.3 | 0.706 | 0.144 | -3.6 |
| no continuous channel | 51.8% | 0.485 | 0.461 | 71.5 | 0.687 | 0.175 | -5.2 |

*chain-of-structures* ranks first on five of the six metrics and second on the sixth. The ordering of the four baselines is informative in itself: DPLM *<* CFP-Gen *<* MMCP-Gen *<* CodeFP follows how much of the design each method states explicitly before generating. DPLM is conditioned only on the sequence prior and reaches 32.3%. CFP-Gen and MMCP-Gen add functional labels and reach the mid-forties. CodeFP conditions on a learned codebook of functional descriptors and reaches 55.3%. *chain-of-structures* adds an explicit intermediate state rather than a richer label, and improves on CodeFP by **1.7** points of EC3. This margin is small, and we interpret it accordingly: writing the design down is worth approximately as much as the strongest label-conditioned generator, not more.

The structural metrics separate the methods more sharply than the functional ones. pLDDT spans 58.2 to 79.3 and TM spans 0.539 to 0.729, and both order the methods as EC3 does. This is relevant because a functional score can be attained by a sequence that does not fold. Here the methods that place function best also fold best, so the functional gains do not come at the cost of designability.

CodeFP obtains the better MMD (0.116 against 0.127), meaning its outputs lie closer to the natural *k*-mer distribution. Proximity to the natural distribution is not equivalent to correctness for the target reaction, and the two metrics disagree on this comparison. Our ablations below exclude the explanation that explicit position-wise commitments move composition away from the background distribution. We do not have a measurement that identifies the remaining cause.

F1-Micro exceeds F1-Macro for every method (0.533 against 0.513 for *chain-of-structures*), which is expected when many EC classes have few members: micro weights samples and macro weights classes, so the macro score is determined by the long tail. The gap is similar across methods, so it reflects the class distribution of the test set rather than a property of any generator. Macro scores computed on the well-populated classes alone rise for every row and compress the differences between rows, so we report the unrestricted macro score, which is the stricter of the two.

Removing the written chain costs **3.6** points of EC3, and removing the continuous channel costs **5.2**. The second figure is the more informative of the two, because the written rounds are still produced in that variant. What is removed is the path by which structural supervision reaches the reasoner, not the plan the reasoner writes.

Neither ablation reaches the strongest baseline. CodeFP obtains 55.3% in Table 3, above both 53.4% and 51.8%. The 1.7-point margin in Table 3 therefore requires the written rounds and the continuous channel together; removing either places *chain-of-structures* below a label-conditioned generator.

The ordering full *>* no-chain *>* no-continuous holds for EC3, both F1 scores, pLDDT, TM and MMD, with no inversion. The two components therefore do not trade one axis against another, and no variant improves function at the cost of designability. MMD in particular moves toward the natural *k*-mer distribution as components are added (0.175, 0.144, 0.127). Had the MMD deficit in Table 3 been caused by explicit position-wise commitments, the ablated variants would score better on MMD; they score worse.

## 6 Discussions

The comparison in Section 5 establishes that writing a design down as an explicit chain is competitive with the strongest label-conditioned generator, but the **1.7**-point EC3 margin also bounds what a purely forward chain buys. The four rounds commit in one direction: each round conditions on the state the previous round produced and never revisits it. We read the size of that margin as the price of this restriction, and the rest of this section sets out what relaxing it would require.

The four-round chain is a starting point for *longer, adaptive scientific reasoning*. Its schedule and two-expert architecture serve this specific case study of *Chain of Science Tokens* and do not define Scinetics’ final model architecture. We aim to let the model inspect a generated state, diagnose a conflict, and decide whether to refine that state or revisit an earlier decision. An incompatible scaffold, for example, could trigger a return to active-site organization; persistent sequence–structure inconsistency could trigger local repacking or backbone revision. Backtracking would explicitly reopen affected commitments before regenerating the dependent states. This extends the current forward schedule to trajectories that include correction, alternative hypotheses, and repeated evaluation.

Expert demonstrations and LLM agent workflows can be extended to supply successful repairs and failed attempts alongside forward designs. Learning when to continue, revise, or stop would connect longer reasoning to measurable improvements under a computational budget. Matched baseline comparisons, sequence-cluster holdouts, and experimental measurements of turnover, selectivity, expression, and stability will guide this development. We ultimately aim to extend Chain of Science Tokens across scientific modalities, with generated objects and experimental feedback shaping each subsequent reasoning step.

